# JACKIE: Fast enumeration of genomic single- and multi-copy target sites and their off-targets for CRISPR and other engineered nuclease systems

**DOI:** 10.1101/2020.02.27.968933

**Authors:** Jacqueline Jufen Zhu, Albert Wu Cheng

**Affiliations:** School of Biological and Health Systems Engineering, Arizona State University, Tempe, AZ 85281, USA; The Jackson Laboratory for Genomic Medicine, Farmington, CT 06032, USA; The Jackson Laboratory Cancer Center, Bar Harbor, ME 04609, USA; Department of Genetics and Genome Sciences, University of Connecticut Health Center, Farmington, CT 06030, USA; Institute for Systems Genomics, University of Connecticut Health Center, Farmington, CT 06030, USA

## Abstract

ZFP-, TALE-, and CRISPR-based methods for genome, epigenome editing and imaging have provided powerful tools to interrogate functions of genomes. Targeting sequence design is vital to the success of these experiments. While existing design software mainly focus on designing target sequence for specific elements, we report here the implementation of JACKIE (Jackie and Albert’s Comprehensive K-mer Instances Enumerator), a suite of software for enumerating all single- and multi-copy sites in the genome that can be incorporated for genome-scale designs as well as loaded onto genome browsers alongside other tracks for convenient web-based graphic-user-interface (GUI)-enabled design. We also implement fast algorithms to identify sequence neighborhoods or off-target counts of targeting sequences so that designs with low probability of off-target can be identified among millions of design sequences in reasonable time. We demonstrate the application of JACKIE-designed CRISPR site clusters for genome imaging.

## INTRODUCTION

Zinc finger protein (ZFP), Transcription Activator Like Effector (TALE) and CRISPR-Cas system have revolutionized genome biology and synthetic biology by providing versatile DNA and RNA-targeting modalities for engineering [1-3]. These technologies have revolutionized genome research by providing versatile tools for “writing” the genome, epigenome and transcriptome as well as “reading” the dynamics of the genome architecture through live-cell genome imaging. The success of these experiments depends heavily on the design of target sequences within the genome. While genome and epigenome editing experiments usually requires target sequences that occur only once in the genome, genome imaging require clustered repetitive sequences [4-7]. Most of the existing design software packages provide small scale or one-by-one design [8-11]. For large-scale design, these software packages are inefficient. Furthermore, for genome imaging where clustered repetitive sequences are needed, no design software is available.

For enumeration of all potential target sites of length k (k-mers) and their locations in the genome, we implemented JACKIE (Jackie and Albert’s Comprehensive K-mer Instances Enumerator) software package that is compatible with high performance computing (HPC) clusters. This allow the generation of databases of single- or multi-copy target sites that can not only be used in genome-wide library design but can also be loaded as genome tracks on genome browsers [12] alongside other genomic tracks to allow for GUI-enabled design of targeting experiments. To demonstrate application, we generated databases of CRISPR target sites in human hg38 and mouse mm10 genomes and selected several clustered gRNAs for genomic imaging experiments.

## MATERIALS AND METHODS

### Implementation of JACKIE

JACKIE consists of four sets of programs implemented in C++, `JACKIE.bin`, `JACKIE.sortToBed`, (`JACKIE.encodeSeqSpace, JACKIE.encodeSeqSpaceNGG, JACKIE.encodeSeqSpace.prefixed, JACKIE.encodeSeqCountDatabase`) and (`JACKIE.countSeqNeighbors, JACKIE.countSeqNeighbors.pmulti, JACKIE.countOffSites`), as well as Python and Bash scripts for job batching and downstream processing (Fig 1). `JACKIE.bin` scans the genome and enumerates any potential targeting sites (k-mers) on both strands (Fig 1a). A pattern-matching parameter allows the specification of length k of the k-mer binding sites, as well as motif constraints (such as NGG in the case of CRISPR-SpCas9). To enhance memory and computation efficiency, k-mers are mapped to an unsigned 64-bit integer (`uint64_t NucKey`) by encoding each nucleotide and an end-of-string (EOS) indicator in 3 bits, A->000, C->001, T->010, G->011 and EOS->111 (Fig 1a). The chromosome (unsigned 32-bit integer, `uint32_t chrID`) and location for each sequence instance (signed 32-bit integer, `int32_t pos`) are recorded such that chID maps to a chromosome through a tab-delimited table while the sign of `pos` registers the positive (+) and negative (-) strands. The aggregate `KeyedPosition` data (`NucKey, ChrID, pos`) generated at each position during the genome scan are written into files prefixed by a predefined prefix length (default to 6, e.g., ATGAGC.bin contains all `KeyedPosition` of k-mers prefixed by ATGAGC) so that the next steps can be highly parallelized. The `JACKIE.bin` operations can be divided into four subtasks each working on k-mers prefixed by each of the four bases (e.g., the “A” subtask generates all AXXXXX.bin files and an A.ref.txt chromosome table). Upon the completion of the `JACKIE.bin` operations, `JACKIE.sortToBed` is run on each bin file by loading the `KeyedPosition` records into a list in memory, and sorting them by `NucKey`, thus placing `KeyedPosition` records of identical k-mers adjacent to each other in the list (Fig 1b). The `NucKey`-sorted list is then traversed to output k-mer sites in bed interval files that can be concatenated to generate a genome-wide target site database (JACKIEdb) (Fig 1c). The extensive divide-and-conquer design and binary encoding of sequences allow `JACKIE` to be highly parallelized on an HPC cluster as well as efficient sorting of sequences. A python script (`chainExonBedsToTranscriptBed.py`) can be used to take bed file outputs from `JACKIE` to further collapse same-chromosome target sites into an extended bed interval file format with each record containing locations of sites with the same k-mer encoded as bed file blocks to appear as chains in genome browsers (see Fig 2b). Other helper scripts (`fastjoinBedByOverlap.py`, `joinBedItemsWithinRadius.py`) can be used for downstream processing, such as for the fast extraction of target sites overlapping defined set of intervals. target sequences, number of occurrences, and span sizes (the distance between the 5’ most site to 3’ most site on the same chromosome) are encoded in the output files so that target sites can be filtered by simple text processing software (e.g., awk) to identify sites or site clusters with desired properties.

**Fig 1.**
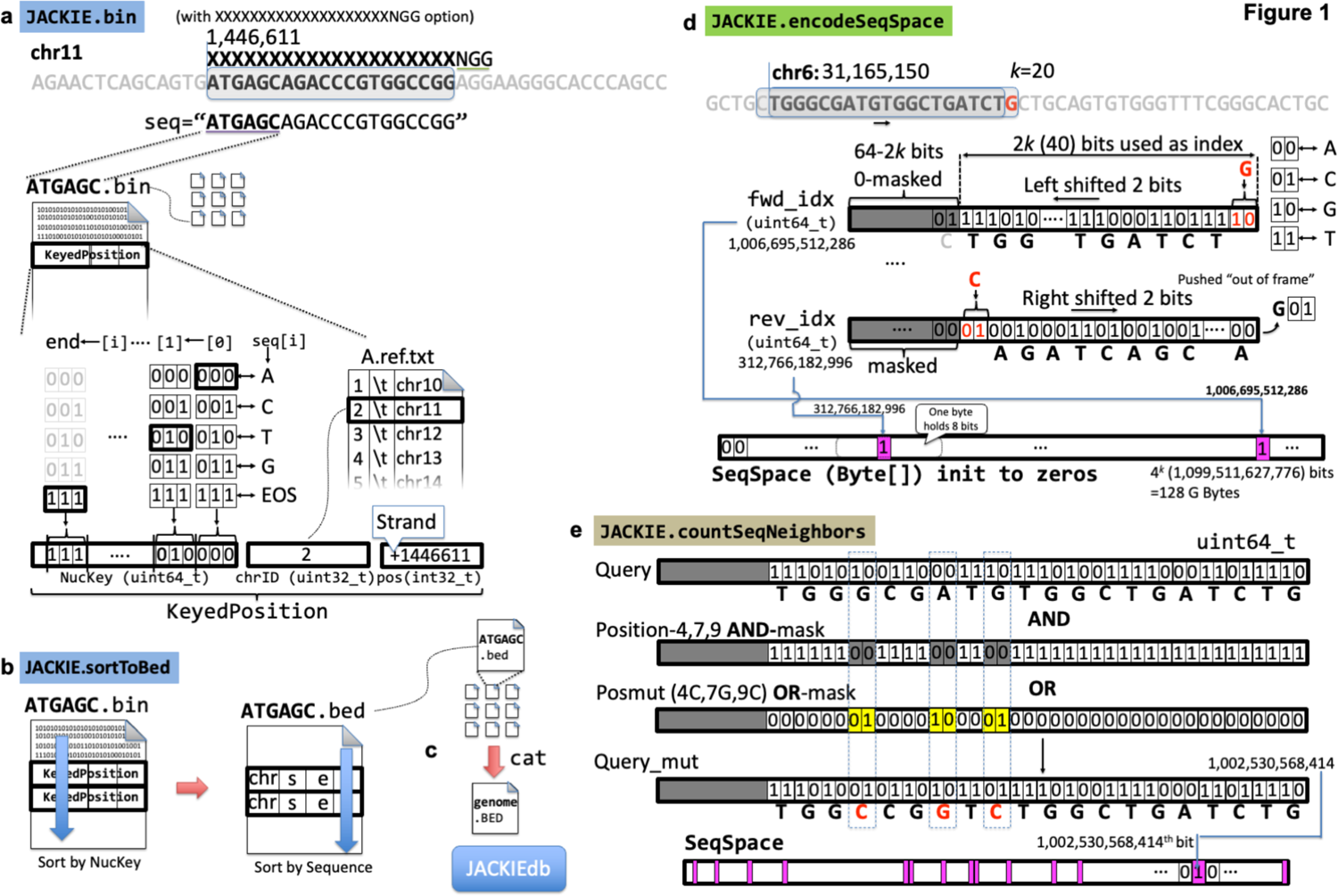
Implementation of JACKIE. **(a)** `JACKIE.bin` scans the genome for k-mers matching a specified motif (e.g., XXXXXXXXXXXXXXXXXXXXNGG matches 20-mer denoted by 20X followed by NGG proto-adjacent motif (PAM) required by the CRISPR/SpCas9 system), and encode each occurrence in a data structure called `KeyedPosition`, consisting of NucKey (unsigned 64-bit integer, `uint64_t`), `chrID` (unsigned 32-bit integer, `uint32_t`), and `pos` (signed 32-bit integer, `int32_t`). NucKey records a binary representation of the k-mer sequence using 3 bits per nucleotide. `chrID` records an integer representing a chromosome. A `chrID`-chromosome mapping file is generated. pos records the location of the binding site, with negative and positive integers representing the minus strand and positive strand, respectively. `JACKIE.bin` outputs `KeyedPosition` to files as it scans the genome. To parallelize this step, four processes are started as separate cluster jobs, each focusing on the A, C, T, or G as the first nucleotide of k-mer. To allow for parallelization in subsequent steps, `KeyedPosition` records are appended to files per 6-mer prefix of the k-mer (<6merPrefix>.bin, e.g., ATGAGC.bin contains all records for all k-mer starting with ATGAGC). **(b)** `JACKIE.sortToBed` loads each <6merPrefix>.bin file and performs a sort on the `KeyedPosition` records on the `NucKey` variable and then traverse the sorted list of `KeyedPosition` to output a bed interval file containing the coordinate position of each k-mer site with item named by <NucKey>.<copy number>/<sequence>, and score field recording the copy number. Parallelization is achieved by either starting separate jobs on individual <6merPrefix>.bin files, or by starting batch jobs on 2mer prefices (e.g., AA AT AC AG operating on AA*.bin, AT*.bin, AC*.bin, AG*.bin, etc). **(c)** The outputs (<6merPrefix>.bed) are then merged into a combined bed file (JACKIEdb) using cat Unix command. **(d)** Evaluation of off-target effects starts with generating a bit array representation of sequence space (`SeqSpace`) of the target genome. `JACKIE.encodeSeqSpace` initializes a 4^*k*^-bit (i.e., 4^2*k*−3^ bytes) array to zeros as an “empty” `SeqSpace`. The program scans the genome and record the presence of each encountered k-mer by first translating the sequence to a binary index, and then setting the bit on the `SeqSpace` referenced by the index to one. The reverse-complement k-mer is handled similarly. A sliding mechanism that shifts two bits as the genome advance to the next nucleotide is used for both indices for time efficiency. **(e)** `JACKIE.countSeqNeighbors` reads in `SeqSpace` and a table of query sequences (e.g., target site design bed files from `JACKIE.sortToBed`) and compute the numbers of sequence neighbors with different number of mismatches up to a specified threshold. At start, `JACKIE.countSeqNeighbors` builds a collection of position AND-masks and pos_mut OR-masks that when applied to a query sequence encoded in the binary representation generates all possible neighbor sequences up to the specified mismatch number. The resultant query_mut values then serve as the indices to query the `SeqSpace` array. If the corresponding bit of the array is 1, the sequence neighbor count for that particular mismatch number is incremented by one. After going through all pairs of AND-mask and OR-mask, the sequence neighbor counts for each mismatch number up to the specified threshold are reported for that particular query sequence.

**Fig 2.**
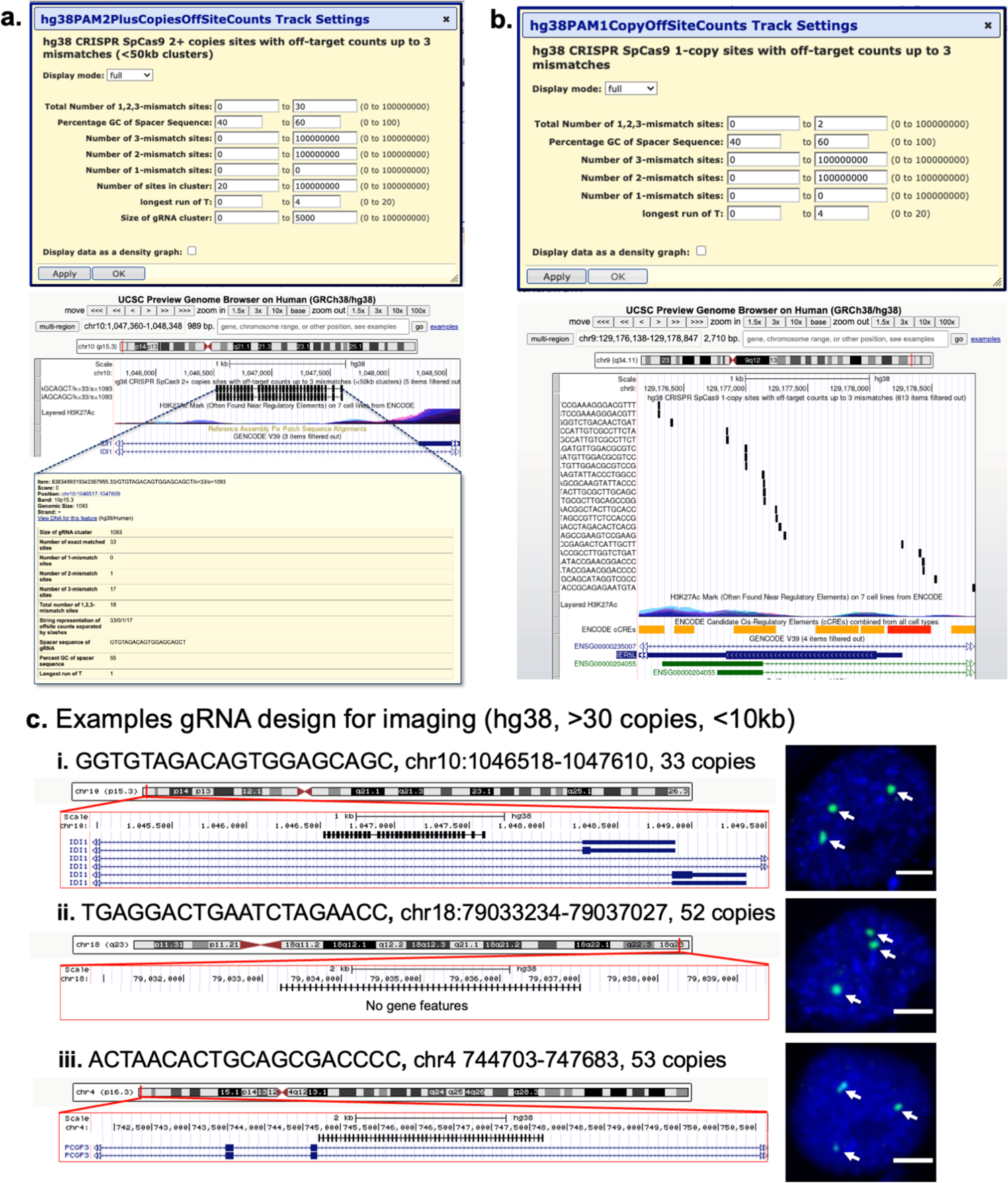
Example use cases for JACKIE designs. **(a)** A screenshot of UCSC genome browser showing CRISPR/SpCas9 gRNA clusters. Top: Example track filter settings for 30 or fewer total off-targets up to 3-mismatches, 40-60%GC spacer, no 1-mismatch off-targets, 20 or more sites in cluster, spacer not containing more than runs of 4 or more T, cluster smaller than or equal to 5kb. Middle: Example gRNA clusters after filtering. Bottom: Information page of the first cluster shown after clicking on the item. **(b)** screenshot of UCSC genome browser showing single-copy CRISPR-SpCas9 gRNA sites. Top: Example track filter settings for up to 2 off-target sites, 40-60%GC spacer, no 1-mismatch off-targets, spacer not containing more than runs of 4 or more T. **(c)** To demonstrate genomic imaging use case, we selected three gRNA clusters and performed CRISPR genome imaging experiment. UCSC genome browser screenshots are shown on the left with the line-bar annotating the CRISPR sites, and representative fluorescence microscopy images shown on the right. Microscopy images were derived by merging green (CRISPR) and blue (DAPI: nucleus stain). Arrows point to fluorescent loci of interest. Scale bars, 5μm.

Off-target effects of engineered nucleases happen when non-target sites with similar sequence to the target sites are recognized and whereat unwanted effects are induced. To prevent or reduce off-target effects, the simplest method is to filter for target sites with fewer predicted off-target locations having same or similar sequences. One of the most popular software for CRISPR off-target predictions is Cas-OFFinder which enumerates sequences and their occurrences with up to a certain number of mismatches to the designed gRNAs [9]. While very efficient in identifying off-target sites for a few gRNAs, evaluation of millions of gRNA designs using Cas-OFFinder would be impractical. In many cases, the goal in identifying a specific target sequence is not about the exact locations of the off-target sites, but the number of off-target sites with a different number of mismatches. And in the case of genome imaging and epigenome editing where there is a relatively higher degree of freedom for target site selection, i.e., the target sites only need to be at close proximity of the element of interest or within a relatively large genomic region, the problem then becomes finding target sites with few or no off-target sequences with up to a certain number of mismatches. Motivated by this, we implement off-target evaluation strategy in terms of “neighborhood” sequences of target sites in the sequence space. The goal is to identify the number of k-mers sequences (in terms of their presence or absence but not their actual locations nor copy numbers) that are certain number of mismatches (e.g., 3), or hamming distances, away from the query design sequence. The off-target prediction routines consist of two programs: `JACKIE.encodeSeqSpace` (or `JACKIE.encodeSeqSpaceNGG` for CRISPR-SpCas9-PAM constrained sequence subspace) and `JACKIE.countSeqNeighbors` (Fig 1d,e). `JACKIE.encodeSeqSpace` scans genome sequence files and record the presence or absence of each possible k-mer in a bit array representation of the k-mer sequence space (SeqSpace) (Fig 1d). At the beginning, SeqSpace is sized at 4^*k*^ bits, i.e., 4^*k*^/8=2^2*k*−3^ bytes, and initialized with zeros. As the genome sequence is scanned, each encountered k-mer sequence is mapped per nucleotide to 2-bit representation (A→00, C→01, G→10, T→11), and packed into a unsigned 64-bit integer (`uint64_t`). The unused most significant bits are zero-masked. The forward strand sequence is mapped to the `fwd_idx` variable while the reverse complementary sequence is mapped to the `rev_idx` variable. To ensure computational efficiency, a “sliding” scheme is used such that the forward index (`fwd_idx`) is shifted left by 2 bits while the reverse complement index (`rev_idx`) is shifted right by 2 bits as the scanning process advances to the next nucleotide, and two bits for the new encountered nucleotide (or the reverse complement) are then added to the least or most significant bit positions, respectively, for `fwd_idx` and `rev_idx`. The bits of SeqSpace at the positions indexed by `fwd_idx` and `rev_idx` is then set to 1, recording the presence of the k-mer and its reverse complement in the SeqSpace. At completion, SeqSpace will have recorded in each bit the presence (bit value 1) or absence (bit value 0) of a corresponding indexed k-mer in the genome. The SeqSpace array will then be outputted to disk as compressed `seqbits.gz` to allow for future uses. `JACKIE.countSeqNeighbors` loads SeqSpace from the `seqbits.gz` file and a list of target site sequences such as the ones in JACKIEdb, and count the number of sequence neighbors each target site sequence has for each number of mismatches up to a user-provided threshold (e.g., 3). The binary representation of k-mers and the indexed SeqSpace allow fast computation of sequence neighbors. `JACKIE.countSeqNeighbors` first generates all position bitwise AND-masks and OR-masks that allow all permutation of sequence changes of a query k-mer through a bitwise AND and a bitwise OR operation, respectively. An example pair of AND- and OR-mask for changing three bases of the query in the binary representation is shown in Fig 1e. For each query sequence, `JACKIE.countSeqNeighbors` applies the AND,OR-mask pairs one-by-one, and lookup the bit indexed by the resultant `Query_mut` variable in the `SeqSpace` array, incrementing the count of sequence neighbors for the corresponding number of mismatches by one. After completing all searches for the query sequence, the number of sequence neighbors is outputted per mismatch number. For computers that do not support 128GB memory for 20-mer `SeqSpace`, we implemented “divide-and-conquer” versions, `JACKIE.encodeSeqSpace.prefixed` and `JACKIE.countSeqNeighbors.pmulti` which divide the problems into prefixed subspaces. For example, using 2 nucleotide (nt) prefix, we divided the problem into 16 subspaces (AA, AC, AG, AT, …), each requiring only 8GB memory for k=20. If needed, the programs allow further division of subspaces. For users interested to obtain the exact off-target site counts, we implemented `JACKIE.encodeSeqCountDatabase` and `JACKIE.countOffSites` which encode SeqSpace with exact copy numbers and report exact off-target site counts, respectively. A user-specified number of bits (*b*) are used to record the copy number of a particular k-mer in the genome up to 2^*b*^−2 copies. If the copy number exceeds 2^*b*^−2, the SeqSpace record is set to 2^*b*^−1, and the copy number is recorded in a “overflow” map in the C++ Standard Template Library (STL) map class, by setting the key to the 2-bit representation of the k-mer (uint64_t) and the value to the copy number.

### CRISPR imaging experiments

gRNA spacer sequences (as listed in Fig 2c) were cloned into gRNA-15xPBSc expression vector via an oligo-annealing protocol as previously described [4]. HEK293T cells were cultivated in Dulbecco’s modified Eagle’s medium (DMEM) (Sigma) with 10% fetal bovine serum (FBS)(Lonza), 4% Glutamax (Gibco), 1% Sodium Pyruvate (Gibco) and penicillin-streptomycin (Gibco). Incubator conditions were 37 °C and 5% CO2. Cells were seeded into 24-well plates the day before being transfected with 50ng pAC1445-dCas9 plasmid (Addgene #73169), 25ng pAC1447-Clover-PUFc plasmid (Addgene #73689) and 250ng sgRNA-15xPBSc plasmid. Cells were fixed, mounted on slides, and imaged 48 hours after transfection on a Leica SP8 confocal microscope.

## RESULTS

We ran `JACKIE.bin` (4 jobs) and `JACKIE.sortToBed` (16 jobs) on a Unix HPC cluster with Intel Xeon compute cores for hg38 and mm10 mouse genome, which took totals of ~35 minutes and ~22 minutes, respectively. On a MacBook Pro laptop (2021; MacBookPro18,3; 14-inch; Apple M1 Pro chip; 16GB ram; 1TB SSD; MacOS 12.2.1 Monterey), running `JACKIE.bin` on hg38 human genome with 4 processes took ~12 minutes, and `JACKIE.sortToBed` with 16 processes, took ~27 minutes. Thus, `JACKIE.bin` and `JACKIE.sortToBed` ran on the MacBook Pro laptop for a total of ~40 minutes. `JACKIE.encodeSeqSpaceNGG` on hg38 took 45 minutes (25 minutes spent on encoding, 20 minutes spent on compressing to seqbits.gz file) on one node of the Unix HPC cluster. We ran `JACKIE.countSeqNeighbors` on 52,666,035 single-copy CRISPR query sequences for up to 3 mismatches which took 12.5 hours on one node of the Unix HPC cluster. On a MacBook Pro laptop (2021; MacBookPro18,3; 14-inch; Apple M1 Pro chip; 16GB ram; 1TB SSD; MacOS 12.2.1 Monterey), running `JACKIE.encodeSeqCountDatabase` on hg38 with 3-bit SeqSpace took 4645 seconds (~1.3 hours) total when 16 runs of 2-nt subspaces (k=20) were executed in serial. On the same machine, running `JACKIE.encodeSeqSpace.prefixed` took a total of 29 min. Sequence Count Database or SeqSpace encoding (“indexing”) is only needed once for a particular genome for a particular k and PAM. We compared speeds of the off-target prediction portion of JACKIE with `Cas-OFFinder` [9] and `FlashFry` [13]. Cas-OFFinder is one of the first and most popular off-target prediction software while FlashFry was evaluated to be the fastest among existing off-target prediction software packages by a previous paper [14]. Indexing of `FlashFry` database took 2 hours. Running `Cas-OFFinder, FlashFry, JACKIE.countOffSites` and `JACKIE.countSeqNeighbors.pmulti` on 100,000 queries took 26873 seconds (~7.5 hours), 3687 seconds (~1 hour), 657 seconds (~11 minutes), and 295 seconds (~5 minutes), respectively. We ran `JACKIE.countOffSites` and `JACKIE.countSeqNeighbors.pmulti` on 1,000,000 queries which took 6224 seconds (~1.72 hours), 2439 seconds (~40 minutes), respectively. Note these are not completely fair comparisons as Cas-OFFinder reports the locations of off-target sites allowing DNA/RNA bulges in addition to mismatches whereas `FlashFry` and `JACKIE.countOffSites` report the number of off-target sites with mismatches only. The fastest approach taken by `JACKIE.countSeqNeighbors.pmulti` reports the number of neighborhood sequences with mismatches only. For large-scale target design projects only requiring target sites with few or no off-targets up to a certain mismatch threshold, off-target prediction approach taken by JACKIE would suffice and provide a time-effective solution. Bed files generated by JACKIE can be uploaded to genome browser alongside other tracks for convenient GUI-enabled target designs. We generated JACKIE database (JACKIEdb) for CRISPR/SpCas9 hg38 and mm10 and created UCSC genome sessions with gRNA tracks on which users can filter for specific gRNA properties, including the numbers of 1-,2-, and 3-mismatch sites, total off-target site number, GC% of spacer sequence, longest run of T as well as number of sites and size of the gRNA clusters (Fig 2a,b). Not only can users filter for desired gRNA properties, but also can they select gRNAs alongside hundreds of tracks already available on the UCSC genome browser. We selected three CRISPR binding clusters and applied CRISPR genome imaging to visualize them via fluorescence microscopy (Fig 2c), producing three fluorescent foci in each case, consistent with the ploidy of the HEK293T cells used in the experiments.

## DISCUSSION

Advances in engineered nuclease technologies have fueled an explosion of genome, epigenome, transcriptome editing and imaging tools. The easily programmable feature as exemplified by the CRISPR systems allows large-scale genome-wide experiments. While some applications, such as genome and epigenome editing, require target sites that are unique, i.e., with sequence occurring only once in the genome, applications such as imaging requires multiple binding sites that are clustered within a particular region. Existing target design software packages focus on individual designs but are inefficient in large-scale design. In this paper, we described a new software package, JACKIE, that enumerates all single- and multi-copy k-mers in a target genome as well as provides fast evaluation of off-target effects. The off-target prediction algorithm is nearly a hundred fold more time-efficient than the most popular software for off-target prediction, making the off-target prediction of millions of sequences practical. Genome-wide design by JACKIE can be conveniently loaded onto GUI-enabled genome browsers alongside other tracks so that users can design target sequences with respect to annotated elements or epigenomic features. The genome-wide design in bed format also allows high-throughput computational selection of target sites using command line tools. We computed CRISPR-SpCas9 single-copy and multi-copy binding sites (JACKIEdb) that can be directly downloaded or visualized on UCSC genome browser. As a demonstration for the utility of gRNA designs by JACKIE, we selected three <10kb clusters of >30 gRNAs and performed CRISPR genome imaging. We believe that JACKIE and JACKIEdb will facilitate designs of genome-wide target libraries or large-scale engineered nuclease projects.

## DATA AVAILABILITY

Source codes and precomputed tracks for hg38 and mm10 are available for download at http://cheng.bio/JACKIE and https://github.com/albertwcheng/JACKIE2

## ACKNOWLEDGEMENTS

This work has been supported by the National Human Genome Research Institute grant R01HG009900 (to A.W.C). Arizona State University startup grant (to A.W.C.) and Jackson Laboratory internal grants (to A.W.C.).

